# Network perturbation analysis of gene transcriptional profiles reveals protein targets and mechanism of action of drugs and influenza A viral infection

**DOI:** 10.1101/175364

**Authors:** Heeju Noh, Jason E. Shoemaker, Rudiyanto Gunawan

## Abstract

Genome-wide transcriptional profiling provides a global view of cellular state and how this state changes under different treatments (e.g. drugs) or conditions (e.g. healthy and diseased). Here, we present ProTINA (Protein Target Inference by Network Analysis), a network perturbation analysis method for inferring protein targets of compounds from gene transcriptional profiles. ProTINA uses a dynamic model of the cell-type specific protein-gene transcriptional regulation to infer network perturbations from steady state and time-series differential gene expression profiles. A candidate protein target is scored based on the gene network’s dysregulation, including enhancement and attenuation of transcriptional regulatory activity of the protein on its downstream genes, caused by drug treatments. For benchmark datasets from three drug treatment studies, ProTINA was able to provide highly accurate protein target predictions and to reveal the mechanism of action of compounds with high sensitivity and specificity. Further, an application of ProTINA to gene expression profiles of influenza A viral infection led to new insights of the early events in the infection.

## INTRODUCTION

The identification of the molecular targets of pharmacologically relevant compounds is vital for understanding the mechanism of action (MoA) of drugs, as well as for exploring off-target effects.While the definition of a target can be quite arbitrary, the term generally refers to a molecule whose interaction with the compound is connected to the compound’s effects (1). In this study, transcription factors (TFs) and their protein interaction partners represent the target molecules, while differential gene expression profiles represent the effects. Among existing technologies for protein target discovery (e.g., biochemical affinity purification, RNAi knockdown or gene knockout experiments) (2), gene expression profiling has received much recent attention due to its relative ease of implementation as well as the availability of large-scale public databases and well-established experimental protocols and data analytical methods. A complication when using gene expression profiling for target discovery is that the data give only indirect indications of the drug’s action. As illustrated in Fig. 1a, the interaction between a compound and its protein target(s) is expected to result in the differential expression of downstream genes that are regulated by the protein target(s). But, the expression of the protein targets themselves may not – and often do not – change (3). Consequently, target discovery using gene expression profiles requires computational methods to identify the (upstream) targets from the (downstream) effects.

**Figure 1.**
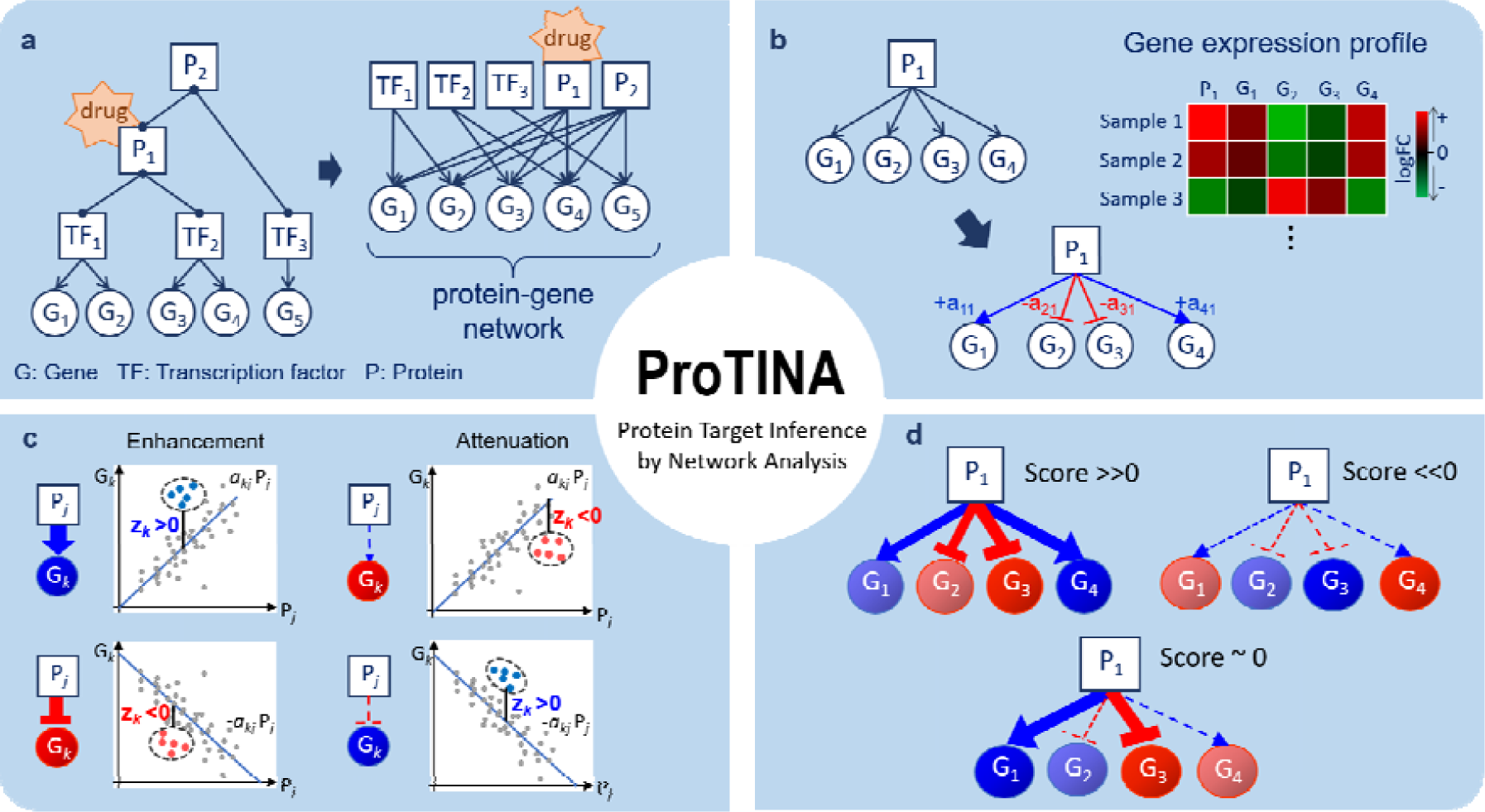
Protein target prediction by ProTINA. (a) The protein-gene network describes direct and indirect regulations of gene expression by transcription factors (TF) and their protein partners (P), respectively. A drug interaction with a protein is expected to cause differential expression of the downstream genes in the PGN. (b) Based on a kinetic model of gene transcriptional process, PROTINA infers the weights of the protein-gene regulatory edges, denoted by *a_kj_*, using gene expression data. The variable *a_kj_* describes the regulation of protein *j* on gene *k*, where the magnitude and sign of *a_kj_* indicate the strength and mode (+*a_kj_*: activation, −*a_kj_*: repression) of the regulatory interaction, respectively. (c) A candidate protein target is scored based on the deviations in the expression of downstream genes from the PGN model prediction (*P_j_*: log2FC expression of protein *j*, G_*k*_: log2FC expression of gene *k*). The colored dots in the plots illustrate the log2FC data of a particular drug treatment, while the lines show the predicted expression of gene *k* by the (linear) PGN model. The variable *z_k_* denotes the z-score of the deviation of the expression of gene *k*from the PGN model prediction. A drug-induced enhancement of protein-gene regulatory interactions is indicated by a positive (negative) *z_k_* in the expression of genes that are activated (repressed) by the protein (i.e. *a_kj_z_k_*> 0). Vice versa, a drug-induced attenuation is indicated by a negative (positive) *z_k_* in the expression of genes that are activated (repressed) by the protein (i.e. *a_kj_z_k_* < 0). (d) The score of a candidate protein target is determined by combining the z-scores of the set of regulatory edges associated with the protein in the PGN. A positive (negative) score indicates a drug-induced enhancement (attenuation). The larger the magnitude of the score, the more consistent is the drug induced perturbations (enhancement/attenuation) on the protein-gene regulatory edges.

Existing computational strategies for compound target identification using gene expression profiles can generally be classified into two groups: comparative analysis and network-based analysis (4). Comparative analysis methods use the gene expression profiles as drug signatures. Here, the similarity between the differential gene expression of a drug treatment and those of reference compounds or experiments with known targets, implies a closeness in the molecular targets and the MoA. A notable example of such an approach is the Connectivity Map (5), which provides gene expression profiles of human cell lines treated by ~5000 small molecule compounds as queryable signatures for evaluating drug-drug similarities (6). The obvious drawback of comparative analysis methods is their dependence on an extensive and accurate target annotation of the reference gene expression profiles.

In network-based analysis, one adopts a system-oriented view by using cellular networks, such as gene regulatory network (GRN) and/or protein-protein interaction network (PIN). A number of network-based analytical methods relied on dynamic models of the GRN to infer network perturbations caused by drug treatments (7-11). Several notable methods include Network Identification by multiple Regression (NIR) (7), Mode of action by Network Identification (MNI) (8), Sparse Simultaneous Equation Model (SSEM) (9) and DeltaNet (10-11). In these methods, the GRN is inferred from a training dataset of gene expression profiles using a linear regression derived from a dynamic mechanistic model of the gene transcriptional process. Subsequently, the inferred GRN is utilized for target identification to evaluate deviations in the differential gene expression caused by drug treatments (7-11) or in disease (12). A major pitfall of the above methods is that the inference of GRN from gene transcriptional profiles is highly challenging (13), as the inference problem often becomes underdetermined (i.e. the GRN may not be inferable) (14-15). In addition, as mentioned above, the expressions of the drug targets are often unaffected by the drug treatment (3).

Another group of network-based analytical methods utilizes cellular network graphs, either curated from the literature knowledge or inferred from gene expression data, to formulate statistical hypothesis tests for ranking drug targets (3, 16-22). Several methods in this category prioritize targets based on the enrichment of the downstream or neighbouring molecules in the network for differentially expressed genes, following the principle of “guilt by association” (3, 16-20). Another set of methods rank targets by scoring hypotheses that are generated based on causal relationships in the biological networks (21-22). A recent method called Detecting Mechanism of Action by Network Dysregulation (DeMAND), combines the GRN and PIN to create a molecular interaction network, where the drug targets are scored based on the statistical significance of drug-induced alterations in the joint gene expression distribution between two connected genes in the network (23). The methods in this group make use of only the (static) topology of cellular networks without much consideration of the dynamics of the gene transcriptional process, and thus are unable to fully exploit information contained in time-series datasets.

In this work, we developed ProTINA (Protein Target Inference by Network Analysis), a network perturbation analysis method for protein target identification using gene transcriptional profiles. The analysis involves two key steps: (a) the creation of a model of tissue or cell type-specific protein-gene regulatory network (PGRN), and (b) the calculation of protein target scores based on the enhancement or attenuation of the protein-gene regulations. In developing ProTINA, we addressed some of the drawbacks in the existing methods. First, the PGRN in ProTINA is based on a dynamic model of the gene transcriptional process, and is therefore able to take advantage of time-series gene expression profiles that are commonly generated by drug treatment studies. In addition, ProTINA leverages on the availability of comprehensive maps of protein-protein and protein-DNA interactions for the construction of the PGRN, which serves as prior information to alleviate network inferability issue. Finally, ProTINA scores candidate targets based on drug-induced perturbations to the expression of genes regulated by the targets, rather than the expression of the targets themselves. We demonstrated the superiority of ProTINA over the state-of-the-art method DeMAND and differential gene expression analysis (DE), in predicting the protein targets of drugs using human and mouse datasets from NCI-DREAM drug synergy challenge (24), genotoxicity study (25) and chromosome drug targeting study (26). Besides protein targets of compounds, we presented the application of ProTINA to study host-pathogen interactions, specifically for elucidating the targets of influenza A viral proteins.

## MATERIAL AND METHODS

### Gene expression data

We applied ProTINA to three datasets of drug treatments from NCI-DREAM drug synergy challenge (24), genotoxicity study (25) and chromosome drug targeting study (26), and to gene expression data of human lung cancer cell Calu-3 from influenza A viral infection studies (27-30). For NCI-DREAM drug synergy challenge, we obtained the raw *Affymetrix Human Genome U219* microarray data from Gene Expression Omnibus (GEO) database (31) (accession number: GSE51068). The raw data were first normalized and transformed into log2-scaled expressions using *justRMA* function in the *affy* package of Bioconductor (32). Then, the log_2_ fold change (log2FC) differential expressions and their statistical significance (Benjamini-Hochberg adjusted *p*-values) were calculated using a linear fit model and empirical Bayes method in the *limma* package of Bionconductor. Three samples from the drug treatment using the low dose of Aclacinomycin A were dropped because all of the log2FC expressions were close to 1 and thus not statistically significant. The probe sets were mapped to gene symbols using *hgu219.db* annotation package (Entrez Gene database as of 27^th^ September 2015). In the case of multiple probe sets mapping to a gene symbol, we assigned the log2FC from the probe set with the smallest average adjusted *p*-value over the samples.

The raw microarray data from genotoxicity study (25) in human HepG2 cell line were obtained from GEO (accession numbers: GSE28878 using *Affymetrix GeneChip Human Genome U133 Plus 2.0* array and GSE58235 using *Affymetrix HT Human Genome U133+ PM* array). As with the drug synergy data, the microarray data were first normalized using *justRMA*, and the log2FCs and their adjusted *p*-values were calculated using *limma* in Bioconductor. Because the data came from different microarray platforms, the gene symbols were matched separately for each platform using *hgu133plus2.db* annotation package (Entrez database of 27^th^ September 2015) and *HT_HG-U133_Plus_PM* annotation file in Affymetrix, respectively. Likewise, in the case of multiple probe sets matching a gene symbol, the probe set with the smallest average adjusted *p*-value across all samples was chosen.

The raw data from the chromosome-targeting study using mouse pancreatic alpha and beta cells (26) were also obtained from GEO database (ascension number: GSE36379). Again, the raw data were normalized using *justRMA*, and the log2FCs and their adjusted *p*-values were calculated by *limma*. The probes were mapped to the corresponding gene symbols using *moe430a.db* package (Entrez Gene database as of 27^th^ September 2015) in Bioconductor. In the case of multiple probe sets mapping to a gene symbol, we selected the probe set with the smallest average adjusted *p*-value among the samples.

For influenza A infection analysis, we obtained the raw microarray data of four influenza studies (27-30) from GEO database (ascension numbers: GSE40844, GSE37571, GSE33142, andGSE28166). The raw data were background-corrected and normalized using *normexp* and *quantile* methods in *limma* package of Bioconductor. The log2FCs and their adjusted *p*-values were again calculated by *limma*. The probes were mapped to the corresponding gene symbols using hgug4112a.db package (Entrez Gene database as of 27th September 2015). Like before, for genes with multiple probe sets, we chose the logFC value corresponding to the probe set with the smallest average adjusted *p*-value.

### Protein target identification using ProTINA

#### Protein-gene network

In ProTINA, the PGRN is a bipartite graph with weighted, directed edges pointing from a protein to a gene (see **Fig. 1a**). The edges in the PGRN describe the regulation of gene expression by TFs and their protein partners, the molecular targets of interest in this work. The bipartite PGRN above is able to capture feedback loops in the gene transcriptional regulation, even though these loops are not drawn explicitly. An example of such a feedback loop is when a protein regulates the expression of its own transcription factor(s). The PGRN is constructed by combining two types of networks, namely the TF-gene network and PIN. For the construction of human cell type-specific PGRNs, we relied on the Regulatory Circuit resource that provides 394 cell type and tissue-specific TF-gene interactions (33). More specifically, for the analysis of the NCI-DREAM drug synergy, genotoxic compound study, and influenza A viral infection study data sets, we used the TF-gene networks of human lymphoma cells, pleomorphic hepatocellular carcinoma cells, and epithelium lung cancer cells, respectively. We included only TF-gene interactions with a Regulatory Circuit confidence score greater than 0.1. The confidence score indicates the normalized promoter activity level in a given cell type (0: not active, 1: maximally active) (33). For the analysis of mouse pancreatic cell dataset, we obtained the mouse pancreatic TF-gene interactions from CellNet (34). In the construction of the PGRNs, any TF-gene interactions involving unmeasured genes were excluded. In summary, the TF-gene network for human lymphoma, hepatocellular carcinoma cell, and epithelium lung cancer cell lines included 31,392 edges pointing from 515 TFs to 5,153 genes, 3,868 edges pointing from 413 TFs to 953 genes, and 42,145 edges pointing from 515 TFs to 7,125 genes, respectively. The mouse pancreatic PGRN included 2,922 edges, involving 95 TFs and 588 genes.

For human PIN, we combined the protein-protein interactions from two databases, namely Enrichr (35) and STRING (36). For mouse pancreatic cells, we obtained mouse (*Mus musculus*) PIN from the STRING database (36). For each TF, we identified its protein partners, defined as proteins that are within a network distance of 2 from the TF in the PIN. When using the STRING database, we included all direct protein partners of TFs, and proteins with a network distance of 2 from TFs with a confidence score reported on STRING larger than 0.5. For human lymphoma, hepatocytes, and lung cancer cells, we identified 11,090 protein partners for a subset of 499 TFs (out of 515 TFs), 10,834 protein partners for a subset of 403 TFs (out of 413 TFs), and 6,175 protein partners for a subset of 504TFs (out of 515 TFs), respectively. For mouse pancreatic cells, we found 6,620 protein partners for a subset of 89 TFs (out of 95 TFs).

Finally, in the construction of the PGRNs, we assigned a directed edge from a TF or from a protein partner of a TF, to every gene regulated by the TF. In summary, the cell type-specific PGRN for human lymphoma cells included 21,488,617 regulatory edges among 11,161 TFs/proteins and 5,153 genes. For hepatocellular carcinoma cells, the PGRN comprised 3,726,393 edges among 10,893 TFs/proteins and 953 genes. For human lung cancer cells, the PGRN comprised 30,656,861 edges among 11346 TFs/proteins and 7,125 genes. For mouse pancreatic cells, the PGRN consisted of 1,417,972 edges among 6,661 TFs/proteins and 588 genes. While increasing the size of the PGRN, for example by including lesser confident TF-gene and protein-protein interactions or by including proteins with a network distance from TFs larger than 2, would allow the scoring of a higher number of proteins, such strategy often lowers the accuracy of the protein target predictions.

#### Gene transcription model

The edges in the PGRN have weights, whose magnitudes represent the strength of the gene regulation and whose signs indicate the direction or the mode of the regulation: positive for gene activation and negative for gene repression. The weights are inferred from the gene expression dataset by adapting a procedure described in our previous method DeltaNet (10-11). The inference of the edge weights is based on an ordinary differential equation (ODE) model of the mRNA production of a gene:

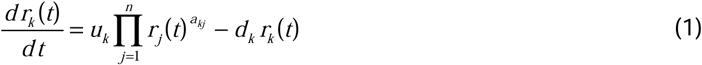

where *r_k_*(*t*) is the mRNA concentration of gene *k* at time *t*, *u_k_* and *d_k_* denotes the mRNA transcription and degradation rate constants respectively, and *a_kj_* denotes the gene regulatory influence (or edge weight) of the *j*-th protein on the *k*-th gene.

While the regulatory edges in the model above usually describe TF-gene interactions, in ProTINA, we further accounted for the (indirect) regulation of a gene by proteins that interact with the TFs. For this purpose, we considered a modified ODE model:

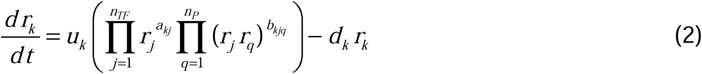

where a positive (negative) *b_kjq_* describes the activation (repression) of the *k*-th gene by a protein *q* through its interaction with the TF protein *j*. The variables *n_TF_* and *n_P_* denote the numbers of TFs and their protein partners, respectively. The multiplication of two variables *r_j_* and *r_q_* implies that the regulation of gene *k* by protein *q* requires the TF protein *j* (a non-zero *r_j_*). The model in Equation (2) can be simplified into:

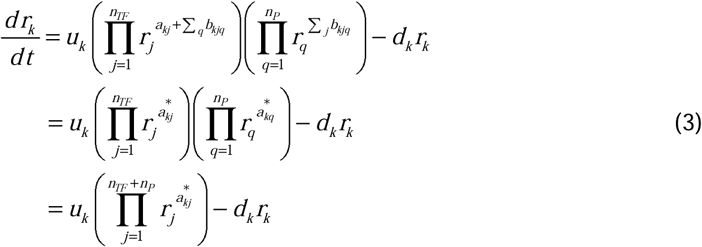

where 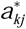 denotes the overall regulatory influence of each protein *j*, including TFs and their protein partners, on the expression of gene *k*. Note that the model in Equation (3) is mathematically equivalent to that in Equation (1).

By taking the pseudo steady-state assumption (i.e. the synthesis rate of mRNA balances the degradation rate, leading to *dr_k_*/*dt* = 0 in Equation (3)), the inference of edge weights 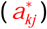 of the PGRN can be rewritten as the following linear regression problem (see derivation in ref. 10):

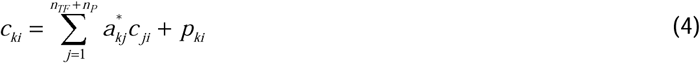

where *c_ki_* denotes the log2 fold-change (log2FC) expression for gene *k* in sample *i*. The variable *p_ki_* represents the part of log2FC of gene *k* expression in sample *i* that cannot be accounted for by the log2FC of its protein regulators. In other words, *p_ki_* indicates the perturbations to the expression of gene *k*. As detailed below, ProTINA relies on the magnitude and directions of such network perturbations (dysregulations) to identify proteins with altered gene regulatory activity.

The dynamical information contained in time-series gene expression profiles could greatly improve the inference of the edge weights above. But, the pseudo steady-state assumption hinders the application of the linear regression in Equation (4) to time-series data. As previously described in ref.11, time-series information could be accounted for by adding the following linear constraint on the linear regression problem:

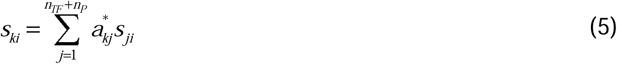

where *s_ki_* is the time derivatives (slope) of the log2FC of gene *k* in sample *i*. In contrast to Equation (4), Equation (5) was derived without assuming pseudo steady-state, which was necessary to account for the dynamics of gene expressions. The slopes of the log2FC at each sampling time point were computed using a second-order accurate finite difference approximation (37). In summary, the estimation of edge weights in ProTINA involved the following linear regression problem:

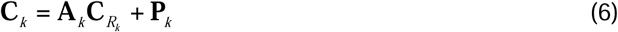

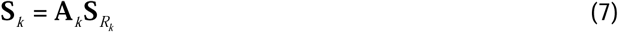

where **C**_*k*_ and **S**_*k*_ are the 1 × *m* vectors of log2FC expressions and time-derivatives of gene *k* across *m* samples, the subscript *R_k_* refers to the set of (*n_TF,k_*+*n_P,k_*) protein regulators of gene *k* in the cell type-specific PGRN, **C***_R__k_* and **S***_R__k_* denote the (*n_TF_*+*n_P,k_*)× *m* matrices of log2FCs and their slopes across *m* samples, **A**_*k*_ is the 1 ×(*n_TF_*+*n_P_*) vector of weights for edges in the PGRN pointing to gene *k*, and **P**_*k*_ is the 1 × *m* vector of dysregulation impacts of gene *k* over *m* samples.

In ProTINA, the vectors **A**_*k*_ and **P**_*k*_ for each gene *k* in Equations (6) and (7) were estimated by ridge regression. The ridge regression provides a solution to an underdetermined linear regression problem of the standard form: **y** = **X***β* + *ε*, using a penalized least square objective function:

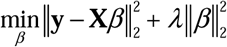

where *λ* is a shrinkage parameter for the L^2^-norm penalty. Equations (6) and (7) are rewritten into the standard linear regression problem with **y** = [**C**_*k*_ **S**_*k*_]^T^, **X** = [ [ **C***_R__k_* **S***_R__k_*]^T^, [*I_m_* **0**]^T^], *β* = [**A**_*k*_ **P**_*k*_]^T^. Before applying the ridge regression, we normalized the vectors of log2FCs and slopes to have a unit norm. Self-loops were excluded in the regression, and thus the diagonal entries of **A**_*k*_ were set to 0. In the applications of ProTINA, we employed 10-fold cross validations to determine the optimal *λ*, one that gives the minimum average prediction error. Here, we used the GLMNET package (38) for both the MATLAB and R versions of ProTINA.

#### Protein target scoring

In ProTINA, each candidate protein target is assigned a score based on the deviation of the expression of its downstream genes. More specifically, we computed the residuals of the linear regression problem in Equations (6) for each gene *k*, i.e.

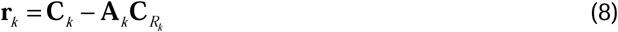

where **r**_*k*_ is the 1 × *m* vector of residuals for *m* samples. For each drug treatment, there often exist multiple gene expression profiles, taken at different time points or different doses. Correspondingly, we evaluated the z-score *z_lk_* for each drug treatment *l* and for each gene *k*, according to

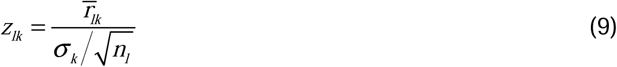

where 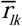 denotes the average residual of gene *k* among the drug treatment samples, *σ_k_* denotes the sample standard deviation of the residuals in all samples besides the drug treatment, and *n_l_* denotes the number of samples from the drug treatment. A positive (negative) z-score indicates that the expression of gene *k* in the particular sample was higher (lower) than expected based on the expression of its regulators. The greater the magnitude of the z-score, the more significant is the gene dysregulation.

The target score of a TF or protein for a drug is calculated by combining the z-scores of the target genes in the PGRN, as follows: (ref. 39)

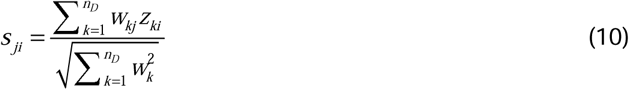

where *z_ki_* denotes the z-score of gene *k* and *s_ji_* denotes the score of the TF/protein *j* in the drug treatment sample *i*. The weighting coefficients *w_kj_* are set equal to the edge weights *a_kj_* divided by the maximum magnitude of *a_kj_* across all *j*. In other words, the weight *w_kj_* reflects the fraction of the regulation of gene *k* expression that could be attributed to protein *j*. When *w_kj_* (or *a_kj_*) and *z_ki_* have the same signs, *w_kj_z_ki_* thus takes a positive value. As illustrated in **Fig. 1c**, a positive *w_kj_z_ki_* implies an enhanced regulatory activity of protein *j* on gene *k*, since the activation (inhibition) of gene *k* expression by protein *j* is stronger in this sample than expected by the PGRN model. In contrast, a negative *w_kj_z_ki_* indicates an attenuation of the regulatory influence of protein *j* on gene *k*, since the activation (inhibition) of gene *k* expression by protein *j* is weaker than predicted by the PGRN model. Consequently, a highly positive (negative) score *s_ji_* is an overall indicator of strongly enhanced (attenuated) regulatory activity of protein *j* by the drug treatment in sample *i* (see **Fig. 1d**). The protein targets in each drug treatment sample are ranked in decreasing magnitude of the scores *s_ji_*.

### DeMAND and differential expression analysis

For DeMAND analysis, we employed the public R subroutines available from the website: http://califano.c2b2.columbia.edu/demand. Following the procedure detailed in the original publication (23), we computed the RMA (Robust Multi-array Average) normalized gene expression values as inputs to the analysis. In DeMAND analysis, we used the same cell type-specific PGRNs as those in ProTINA. For each candidate protein target, DeMAND evaluated the *p*-value of the deviations in the gene expression relationship between the protein target and each of the genes connected to this protein in the PGRN. The drug targets were ranked in increasing magnitude of the combined *p*-values.

In differential expression (DE) analysis, we calculated the log2FC differential expression of each protein in the PGRN, as described in section **Gene expression data** above. Here, we used the log2FC values directly as the target scores. Correspondingly, we ranked the candidate protein targets in decreasing magnitude of the log2FC gene expression values.

### Performance assessment

For comparing the performance of different methods, we computed the area under the receiver operating characteristic curve (AUROC), i.e. the area under the plot of true positive rate against false positive rate, following the procedure adopted in DREAM challenges (40-41). For each method and each drug treatment, we generated a ranked list of protein targets according to decreasing magnitudes of the protein scores in ProTINA, increasing *p*-values of network dysregulation from DeMAND, and increasing magnitudes of log2FC gene expression from DE analysis.

### Gene set enrichment analysis

For influenza A virus study, we performed a gene set enrichment analysis (GSEA) of the protein target predictions from ProTINA, DeMAND and DE analysis for the KEGG biological pathways (42), using the R package GAGE (Generally Applicable Gene-set/pathway Enrichment analysis) with Kolmogorov-Smirnov tests (43). In the case of ProTINA and DeMAND, target proteins with zero score were excluded from the GSEA.

### Reference protein targets

The reference protein targets of compounds in drug treatment studies were compiled from 5 different public databases of chemical-protein interactions: DrugBank (44), Therapeutic Target Database (TTD) (45), MATADOR (46), Comparative Toxicogenomics Database (CTD) (47), and STITCH (48). DrugBank and TTD provided information on the mechanism of drug actions as well as the proteins that have physical binding interactions with drugs. Meanwhile, MATADOR, CTD, and STITCH gave interactions between proteins and chemical compounds, curated from text mining and experimental evidences. When retrieving the protein targets of drugs from these databases, we collected proteins that directly bind to the queried drugs. The reference targets for each dataset in this study are provided in **Supplementary material 1**. Meanwhile, the reference protein targets for influenza A virus study were obtained from ref. 49, where 1,292 host proteins that likely physically bind to viral proteins of influenza type A/WSN/33 in human embryonic kidney cells (HEK293) were identified by whole-genome co-immunoprecipitation assays.

## RESULTS

### New protein target prediction strategy

ProTINA takes advantage of the availability of comprehensive protein-protein and protein-DNA interaction databases to construct, when possible, a tissue or cell type-specific PGRN. The method considers a PGRN with weighted directed edges (see **Fig. 1a**), describing direct and indirect gene transcriptional regulation by TFs and their protein partners. The edge weights are determined by applying ridge regression using the gene expression data based on a kinetic model of the gene transcriptional process (see **Fig. 1b** and **Material and Methods**). Here, a positive weight indicates a gene activation, while a negative weight implies a gene repression. Because of the underlying kinetic model, ProTINA is able to incorporate dynamical gene expression data, a common type of data from drug treatment studies (5, 24-26). The scoring of drug targets is based on the enhancement or attenuation of protein-gene regulatory interactions caused by the drug treatment. A drug-induced gene regulatory enhancement occurs when the expression of genes that are positively (negatively) regulated by a candidate target, becomes higher (lower) in drug treated samples than what is predicted by the PGRN model (see **Fig. 1c**). A drug-induced attenuation describes the opposite scenario, where the expression of positively (negatively) regulated genes of a target is lower (higher) than expected from the model. For any given differential gene expression sample, a candidate protein target is scored based on the overall enhancement and/or attenuation of its regulatory influence on the downstream genes (see **Fig. 1d** and **Material and Methods**). Thus, a protein target with a more positive (negative) score is considered a more likely target of the drug, in which the drug treatment enhances (attenuates) the gene regulatory activity.

### Prediction of known targets of drugs

We tested ProTINA’s performance in predicting drug targets using gene expression data from three drug treatment studies employing human and mouse cell lines. The first dataset came from the NCI-DREAM drug synergy study using human diffuse large B cell lymphoma OCI-LY3 (24), the second from the compound genotoxicity study using human liver cancer cells HepG2 (25), and the third from the chromatin-targeting compound study using mouse pancreatic cells (26). We compared ProTINA to the state-of-the-art network-based analytical method DeMAND (23), and to the traditional differential expression analysis (DE). For the analysis of datasets from human cell lines, we constructed cell-type specific PGRNs by combining human PIN from STRING (36) and Enrichr database (35) and human cell-type specific protein-DNA networks from Regulatory Circuit resource (33). Meanwhile, for the construction of mouse pancreatic cell type-specific PGRN, we used mouse (*Mus musculus*) PIN from STRING (36) and mouse protein-DNA interactions from CellNet (34) (see details in **Material and Methods**).

In assessing the performance of ProTINA and the other methods, we compared the ranked list of protein target predictions for each compound with the reference drug targets compiled from the literature (see **Material and Methods** and **Supplementary material 1**). **Fig. 2** (also see **Supplementary Table S1-3**) summarizes the AUROCs of the target predictions from ProTINA, DeMAND, and DE analysis, showing ProTINA significantly outperforming DeMAND and DE analysis for all three datasets. Here, the drug target predictions from DE analysis had the poorest AUROCs with an overall average below 0.66 (AUROC range: 0.393 – 0.982). Meanwhile, the target predictions of DeMAND were slightly better than the DE analysis, averaging at 0.74 (AUROC range: 0.405 – 0.989) for the three datasets. Meanwhile, ProTINA gave the highest average AUROCs among the methods with an average of 0.83 (AUROC range: 0.425 – 0.991).

**Figure 2.**
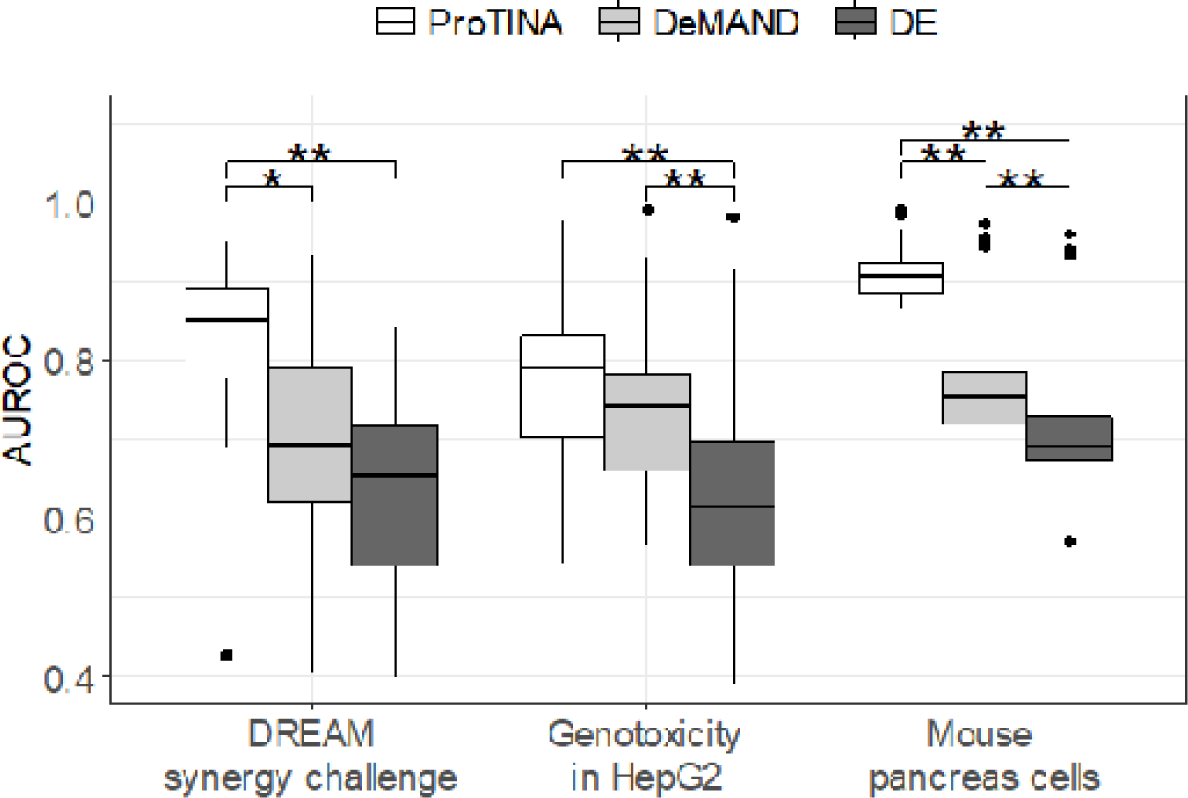
Prediction of known targets of drugs. AUROCs of protein target predictions from ProTINA, DeMAND and DE methods for the NCI-DREAM drug synergy (human B-cell lymphoma), the compound genotoxicity (human HepG2) and the chromatin targeting study (mouse pancreatic cell) datasets (*: p-value < 0.01, **: p-value <0.001 by paired t-test).

### Mechanism of action of drugs

Besides high AUROCs, ProTINA also provided accurate and specific indications on the MoA of the compounds. In the NCI-DREAM synergy study, roughly half of the compounds are known to induce DNA damage response, including DNA topoisomerase inhibitors (camptothecin, doxorubicin and etoposide), DNA crosslinker (mitomycin C), oxidative DNA damaging agent (methothrexate), and histone deacetylase (HDAC) inhibitors (trichostatin A). In demonstrating ProTINA’s ability to reveal the compound MoA, we focused on the canonical p53 DNA damage response pathway (23), as illustrated in **Fig. 3**. Here, the activation of p53 in response to DNA damage is expected to induce the transcription of Cyclin Dependent Kinase Inhibitor 1A (CDKN1A) and Growth Arrest and DNA Damage Inducible Alpha (GADD45A) (50-51). In turn, CDKN1A and GADD45A – through their interactions with Proliferating Cell Nuclear Antigen (PCNA) – regulate the DNA replication and repair process (52). GADD45A also inhibits the catalytic activity of Aurora Kinase A (AURKA) (53), leading to a lowered activation of Polo-like Kinase 1 (PLK1) and Cyclin B1 (CCNB1) in a phosphorylation cascade (54-55). As shown in **Fig. 4a**, except for trichostatin A, the six proteins in the canonical p53 pathway above were ranked highly by ProTINA among the genotoxic compounds in the study (median rank <500), consistent with their known MoA. Note that the same six proteins were ranked much lower among the non-DNA damaging compounds (median rank >500), signifying a high specificity of ProTINA predictions (see also Supplementary Figure S1). Equally important, ProTINA was able to accurately identify the direction of the drug-induced alterations caused by the DNA damaging compounds. The signs of protein target scores from ProTINA indicated drug-induced enhancement (positive scores) of CDKN1A, PCNA, and GADD45A, and attenuation (negative scores) of CCNB1, AURKA, and PLK1 (see **Supplementary Table S4**), consistent with the expected response of these proteins to DNA damage in **Fig. 3**.

**Figure 3.**
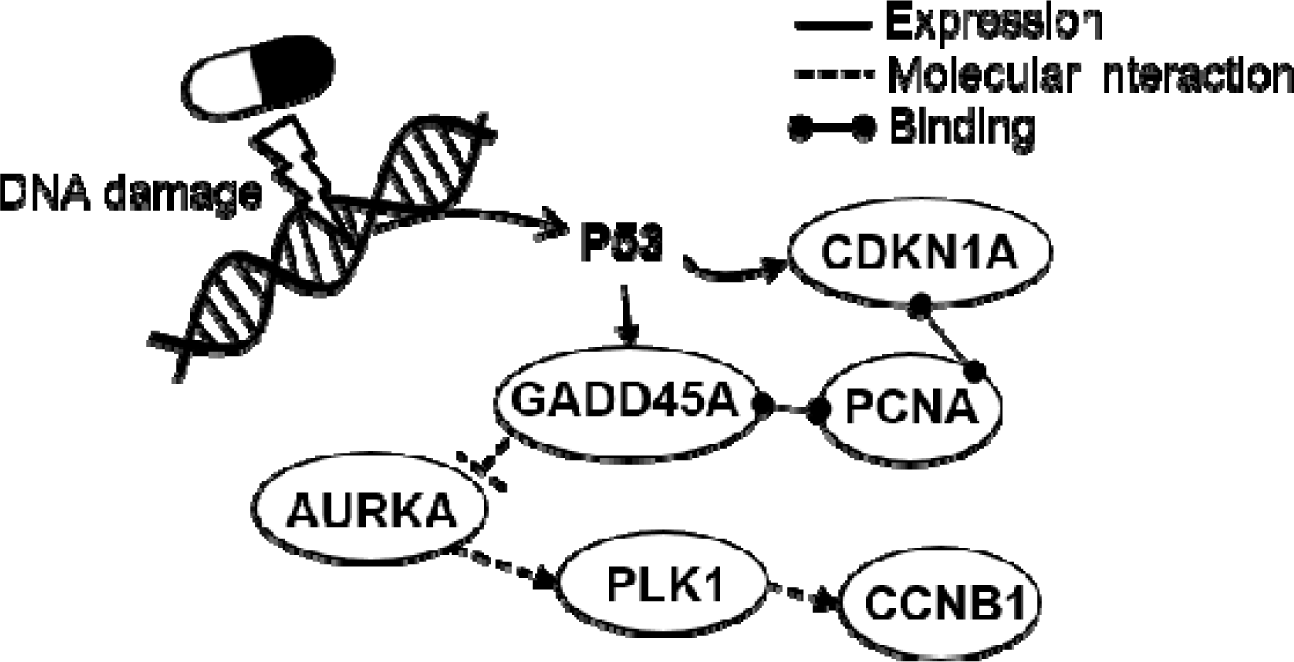
Canonical p53 DNA damage response pathway. In response to DNA damage, GADD45A, CDKN1A, PCNA are activated, while AURKA, CCNB1, and PLK1 proteins are inhibited (23).

**Figure 4.**
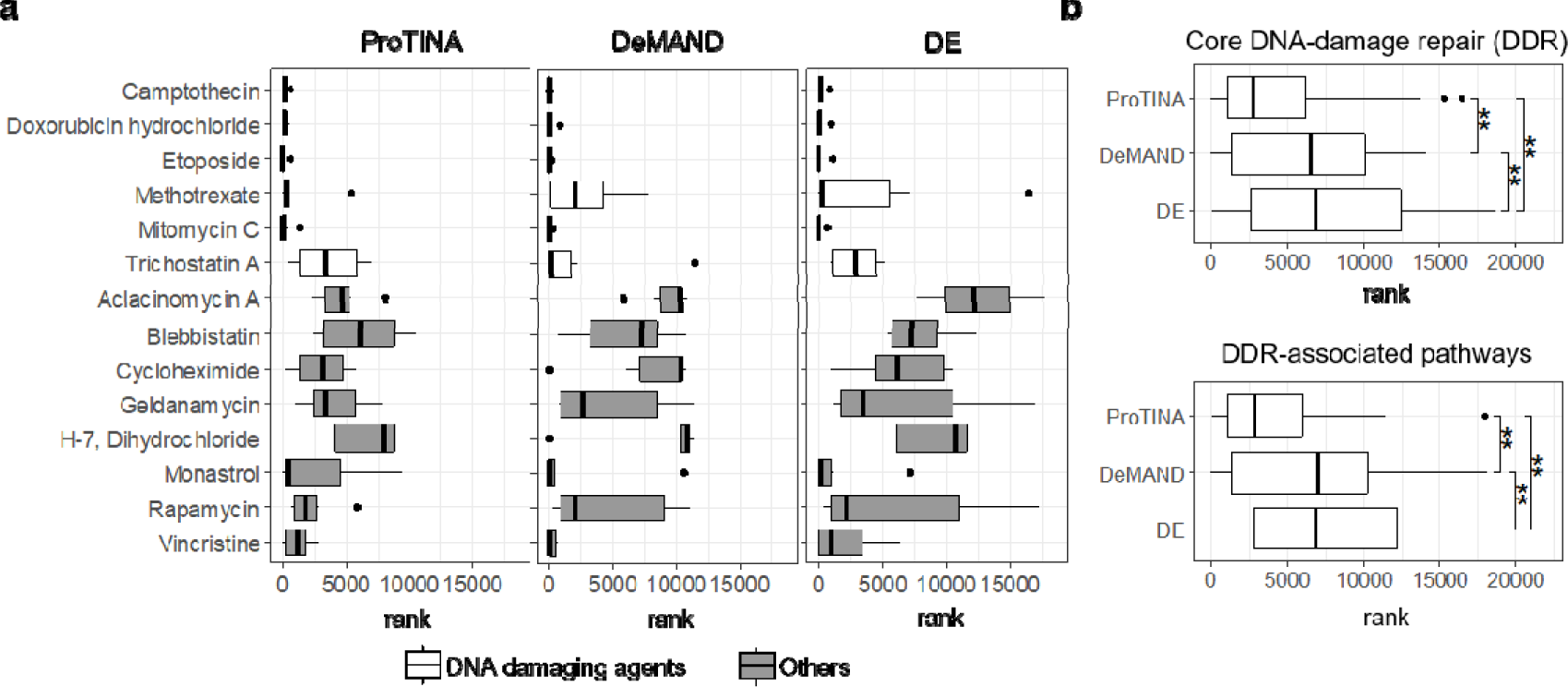
Mechanism of action of compounds based on target predictions by ProTINA. (a) The rank distribution of the canonical p53 DNA damage response proteins in the drug target predictions of PROTINA, DeMAND and DE for the NCI-DREAM drug synergy dataset. (B) The rank distribution of proteins involved in the core DNA-damage repair (DDR) and DDR-associated pathways (56) in the target predictions of PROTINA, DeMAND, and DE for the DNA damaging compounds in the NCI-DREAM drug synergy study (**: p-value <0.001 by Wilcoxon signed rank tests).

As illustrated in **Fig. 4a**, DeMAND and DE analysis also performed reasonably well in predicting the compounds’ MoA. But, the directions of the perturbations predicted by DE analysis were not always consistent with the expected response to DNA damage (see **Supplementary Table S5-6**). Meanwhile, DeMAND did not provide any information on the directions of the drug perturbations. In addition, the protein target scores of ProtTINA provided a clearer demarcation between the genotoxic and the non-genotoxic agents among the compounds in the dataset, than DeMAND and DE analysis (see Supplementary Figure S1). Besides the canonical p53 response pathway, we further looked at the ranking of proteins involved in the overall DNA damage repair (DDR) and its associated pathways (56) (see **Supplementary material 2**). As depicted in **Fig. 4b**, ProTINA ranked these proteins much higher than DeMAND and DE analysis, with DE performing the poorest among the methods considered.

In comparison to DeMAND and DE analysis, ProTINA was further able to detect a specific MoA of mitomycin C, whose DNA crosslinking activity is expected to prompt a particular DNA repair process called the fanconi anemia pathway (57). The fanconi anemia pathway relies on a specific protein complex to ubiquitinate Fanconi Anemia Group D2 Protein (FANCD2) and Fanconi Anemia Group I Protein (FANCI), as well as two homologous recombination (HR) repair proteins, namely Breast Cancer Type 1 Susceptibility Protein (BRCA1) and RAD51 Recombinase (RAD51) (58). In ProTINA analysis, the average rank of FANCD2, FANCI, BRCA1, and RAD51 was within top 100 for mitomycin C, while the average rank of those proteins was much greater than 100 for the other DNA damaging agents (see **Supplementary Table S7**). However, the specific activation of the fanconi anemia pathway by mitomycin C was not detected by DeMAND or DE analysis. Thus, ProTINA provided more sensitive and specific indications for the mechanism of action of compounds than DeMAND and DE.

### Application of ProTINA for predicting pathogen-host interactions

We applied ProTINA to time-course gene expression profiles of human lung cancer cells (Calu-3) under influenza A virus infection, with the goal of identifying host factors that interact with the viral proteins. The gene expression data came from four studies of influenza A viruses, including A/Netherlands/602/2009 (H1N1), A/CA/04/2009 (H1N1), and A/Vietnam/1203/2004 (H5N1) (27-30). We employed ProTINA to compute the overall protein target scores using the gene expression data of Calu-3 from the four studies above, by averaging the scores from the early phase of the influenza infection between 0 to 12 hours. We checked the target predictions of ProTINA against the findings from a genome-wide co-immunoprecipitation analysis of host and viral protein interactions (49). More specifically, the aforementioned study reported 1,292 host proteins that co-immunoprecipitated with viral proteins of influenza A/WSN/33 using human embryonic kidney cells (HEK293). Despite the discrepancy in the cell types and influenza viral strains between the co-immunoprecipitation analysis and the gene expression profiling, influenza A viruses share similar features and common protein interactions (59-60). Besides ProTINA, we also evaluated the accuracy of viral target predictions from DeMAND and DE for the same dataset.

**Fig. 5** gives the receiver operating characteristic (ROC) curves of the target predictions from ProTINA, DeMAND and DE analysis. ProTINA outperformed the two other methods, providing the highest AUROC (ProTINA: 0.76 vs. Demand: 0.69 and DE: 0.65). We further performed a gene set enrichment analysis (GSEA) for the target predictions from each of the methods (see **Material and Methods**) to elucidate the key pathways involved in the viral infection and the accompanying host response. The results of the GSEA are summarized in **Fig. 6**. Both DeMAND and DE target predictions were enriched for only a few pathways (*q*-value < 0.01), while ProTINA prediction had a much higher number of overrepresented pathways.

**Figure 5.**
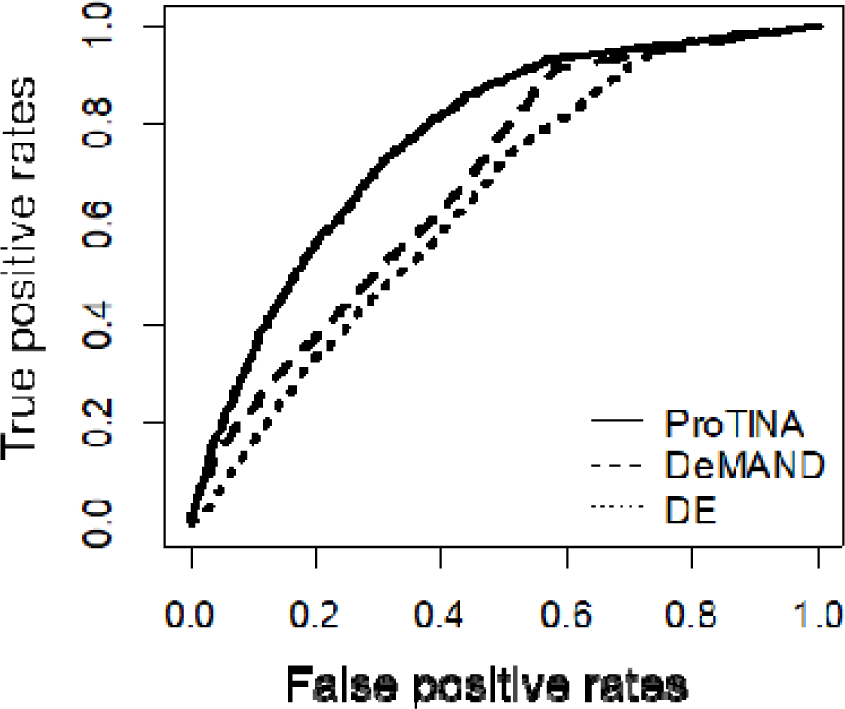
Prediction of targets of influenza A virus. The receiver operative characteristic curves give the true positive rate versus the false positive rate relationship of the protein target predictions from ProTINA, DeMAND, and DE against proteins that co-immunoprecipitate with influenza A viral proteins. The AUROCs for ProTINA, DeMAND and DE analysis are 0.77, 0.69 and 0.65, respectively.

**Figure 6.**
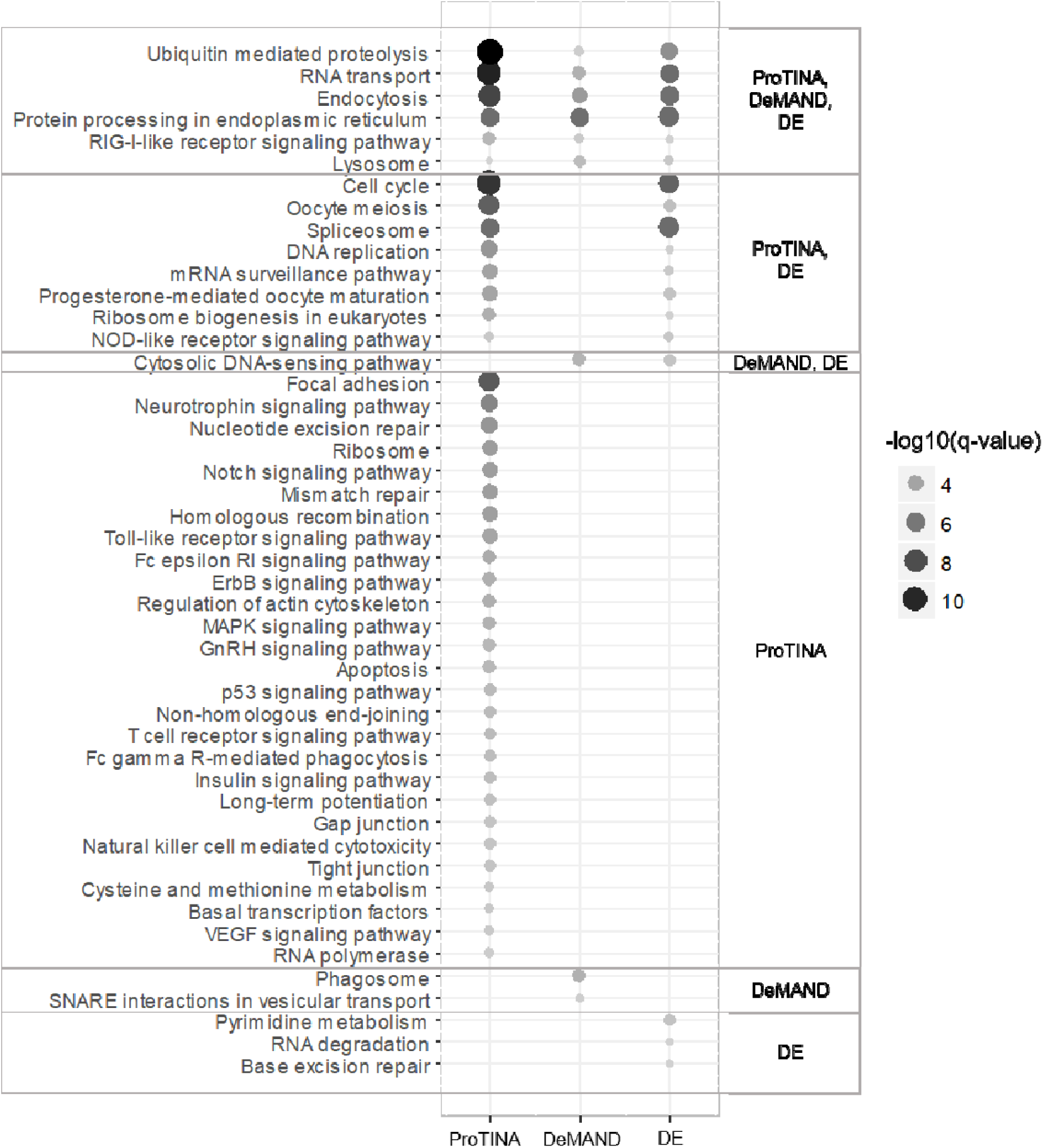
Gene set enrichment analysis for KEGG pathways for the influenza A protein target predictions from ProTINA, DeMAND, and DE. The size of the circles corresponds to −log 10 scale of the *q*-values. Only pathways with *q*-value < 0.01 are shown.

The common enriched pathways among ProTINA, DeMAND and DE (top of **Fig. 6**) included known mechanisms related to viral entry, replication and assembly, including endocytosis (61), protein processing in endoplasmic reticulum (62), ubiquitin mediated proteolysis (63-64) and RIG-I-like receptor signaling pathway (65-66). Both ProTINA and DE analysis indicated the modulation of host cell cycle (67), mRNA surveillance (68) and DNA damage response (69). Only ProTINA prediction was significantly enriched for focal adhesion and actin cytoskeleton, which have been shown to regulate influenza virus entry at the early stage of infection (70). In addition, ProTINA target predictions were also enriched for a broad set of host response pathways to viral infection, including host defense mechanism (e.g., T- and B-cell receptor pathways, phagocytosis, leukocyte migration, chemokine signaling pathways), DNA damage repair (e.g., nucleotide excision repair, p53 signaling pathway, homologous recombination) and apoptosis. As several influenza proteins are known to interfere with interferon production (which in turn regulates several cytokines) (65-66), these findings suggest that, overall, ProTINA provided a broader picture of the early events in the influenza A viral infection, than DeMAND and DE analysis.

## DISCUSSION

ProTINA is a novel and highly effective network-based analytical method for inferring the protein targets of compounds from gene expression profiling data. ProTINA combines the information of TF-gene and protein-protein interactions and data of differential gene expressions to create a tissue or cell type-specific PGRN model. Similar to network-based analysis methods such as NIR (7), MNI (8), SSEM (9) and DeltaNet (10), ProTINA uses a dynamic mechanistic model of the gene transcriptional process to compute deviations in the differential gene expression profiles that are induced by drug treatments. However, as mentioned earlier, the expression of the targets of a drug is often unaffected by the drug treatment (3). For this reason and as illustrated in Fig. 1c-d, ProTINA further transforms the deviations in the differential gene expression into alterations in the protein-gene regulatory edges in the PGRN model. Finally, the target scoring is based on edgetic perturbations of the PGRN, specifically enhancement or attenuation of gene regulatory interactions, caused by the compound.

Like ProTINA, the state-of-the-art method DeMAND also relies on gene transcriptional dysregulations to score drug targets. But, DeMAND does not consider the mode nor the dynamics of the gene regulations, and is unable to predict the direction of the drug-induced dysregulations. DeMAND calculates protein dysregulation scores (*p*-values) for a given gene regulatory network, by statistical comparison between samples from drug treatment and from control experiments. Consequently, DeMAND requires only few samples to generate its prediction (provided that the network can be defined a *priori*). On the other hand, ProTINA makes use of the available differential gene expression profiles from a study or a cell line (i.e., not only from the specific drug treatment), to assign the edge weights of the PGRN by ridge regression. Importantly, in the regression analysis, the PGRN model used in ProTINA accounts for the network perturbations. The ability of ProTINA to incorporate data from other drug treatments or conditions in the scoring of protein targets makes this method particularly suited to take advantage of the extensive and still growing number of gene transcriptional profiles from online databases, such as GEO. As demonstrated in the applications to three benchmark drug treatment datasets using human and mouse cell lines, ProTINA significantly outperformed DeMAND and the standard DE analysis. The target predictions of ProTINA also provide indications for the MoA of compounds, including the directions of the network perturbations, with high sensitivity and specificity.

Besides its intended use to predict targets of compounds, we also demonstrated that the analysis of network perturbations using ProTINA could provide insights into the mechanism of diseases. In the application to gene expression profiles of Calu-3 cells from influenza A infection studies, ProTINA again outperformed DeMAND and DE analysis in identifying host factors that bind with viral proteins. Furthermore, the GSEA of ProTINA target predictions revealed the spectrum of cellular processes involved in the early phase of influenza A infection, including pathways involved in viral entry, replication and assembly, and those related to cellular response to viral infection. Among the pathways with the highest significance (lowest *q*-value) was focal adhesion, which has been shown to regulate influenza viral entry as well as viral replication (70). Meanwhile, the target predictions of DeMAND and DE analysis had fewer enriched pathways, and thus were less informative than the target analysis by ProTINA.

The PGRN model (see Equation (1)) belongs to a class of modeling framework called Biochemical Systems Theory, specifically the S-systems model (71). In addition to gene regulatory networks, S-system modeling have also been used to describe other cellular networks, including signal transduction pathways and metabolic reaction networks (72). Therefore, the principle used in ProTINA could be readily adapted to infer perturbations in cellular signalling or metabolic networks, for example from proteomic and metabolomics profiles, respectively. Besides PGRNs and gene transcriptional profiles, we have not applied ProTINA to analyze other types of cellular networks and data, as such an application was beyond the scope of our work.

ProTINA requires a cell type- or tissue-specific PGRN as an input, which may hinder its application to analyze data from lesser studied organisms. In the case studies, we leveraged on the extensive online databases of protein-protein interactions and TF-gene networks to manually curate PGRNs for human and mouse cells (33-34, 36). Alternatively, provided that a large dataset of gene expression profiles are available for the cell of interest, the PGRN could be inferred using existing network inference methods (73-74). Another potential limitation in applying ProTINA is the requirement for differential gene expression data for inferring the edge weights of the PGRN. While the minimum number for implementing ridge regression with a 3-fold cross validation (lowest fold in GLMNET) is three, the accuracy of the weights and thus the target predictions from ProTINA would generally deteriorate with lower sample sizes. Nevertheless, ProTINA was still able to provide reasonably accurate predictions using a total of 18 samples in the influenza A virus case study above.

The performance of ProTINA, like any other network-based analytical methods, depends on the fidelity of the network used in the analysis. Uncertainty in the PGRN model, both in the structure and the edge weights, is expected to negatively affect the accuracy of the target prediction. Here, structural uncertainty is associated with the reliability of the information used to construct the PGRN, which in our study, comes from online databases of PIN and TF-gene networks. On the other hand, the uncertainty in the edge weights is associated with multiple factors, including the information content of the gene transcriptional profiles and the mathematical formulation used for the weight inference. The information content of the gene expression data is in turn related to measurement uncertainty and richness in the experimental perturbations. Keeping the same number of treatments, datasets with more replicates and less correlated gene expression profiles (i.e. the treatments induce more distinct perturbations to the network), would have a higher degree of information. Meanwhile, we have previously shown that the validity of the model assumption (e.g., pseudo steady-state condition) has an effect on the accuracy of the inferred weights and thus the target prediction accuracy (10). While we have circumvented the issue arising from the violation of the pseudo steady-state assumption in ProTINA (see Equation (7)), (in)validating all model assumptions may be difficult, if not impossible, in practice. A common strategy, as implemented in this study, is to test the performance 519 of the method against benchmark datasets (13). The results of applying ProTINA to drug treatment and influenza A viral infection datasets give confidence to the suitability of the mathematical formulation used in this work.

## ACKNOWLEDGEMENT

We would like to thank Ziyi Hua for her assistance in preparing the R codes of PROTINA.

## AVAILABILITY

MATLAB and R versions of ProTINA can be downloaded from Github repository (https://github.com/CABSEL/ProTINA).

## SUPPLEMENTARY DATA

Supplementary Data are available at NAR online.

